# Stacked MXene/PEDOT-PSS Electrode Fiber for High-Performance Recording and Stimulation

**DOI:** 10.1101/2024.08.20.608461

**Authors:** Shuchun Gou, Peixuan Li, Shu Yang, Guoqiang Bi, Zhanhong Du

## Abstract

The development of microelectrodes with high electrical performance is imperative, particularly for invasive interfaces such as deep brain stimulation (DBS) electrodes. MXene, a new family of 2D early transition metal carbides or nitrides, exhibits outstanding electrical properties and has been researched to improve bioelectronic interface. Through a wet spinning process, we fabricate MXene/PEDOT-PSS electrode fibers measuring 30 μm in diameter, exhibiting an electrical conductivity of 2.16 ± 1.46 × 10^5 S/m and notably low interfacial impedance. The excellent cathodic charge storage capacity (CSCc) and charge injection capacity (CIC) lead to their high performance in recording or stimulation. The electrode fibers are electrochemically stable, biocompatible, magnetic resonance imaging (MRI)-compatible, and demonstrate excellent performance in electromyography (EMG), electrocardiograph (ECG), cortical recording and subthalamic nuclei deep brain stimulation (STN-DBS) experiments.

## Main Text

Recent advancements in materials science and electronics have catalyzed the development of bioelectronic technologies used in medical diagnostics, healthcare monitoring, and wearable devices [1, 2]. High-performance electrophysiological interfaces offer more efficient and safer interventions or recordings for muscle functions, cardiac health, and brain activity, thereby enhancing human-machine interactions and the treatment of neurological disorders. To further achieve miniaturization, there is an urgent need to develop microelectrode materials with high electrical performance, especially for invasive neural interfaces [3, 4].

MXene, a new family of 2D early transition metal carbides or nitrides, has been widely researched since its discovery in 2011. MXene exhibits a graphene-like nanostructure with a high specific surface area, hydrophilic surfaces rich in functional groups, and unique metallic conductivity, along with excellent mechanical, electrochemical, photothermal, and physical properties [5-7]. Initially focused on energy storage, its attractive features have gradually led to applications in chemical sensing and biomedical fields such as photothermal therapy [8-10].

Recently, Ti3C2 MXene has been successfully used in neural stimulation and recording, demonstrating significant potential as a bioelectronic interface [11-13].

Although existing studies based on Ti3C2 MXene have improved the performance of electrode arrays, their enhancement is not significantly greater compared to studies involving graphene oxide (GO). This discrepancy does not align with the outstanding electrical properties of Ti3C2 MXene. We found that electrodes made from Ti3C2 MXene, which the nanosheets solely form the conductive network exhibit superior performance [11]. Reflecting on the field of GO research, we also noted that GO fiber electrodes, formed directly by stacking nanosheets, often exhibit higher cathodic charge storage capacity (CSCc) and charge injection capacity (CIC) compared to microelectrodes formed by other methods, which enhances both recording and stimulation capabilities of the electrodes [14, 15]. Thus, we believe that fiber electrodes prepared by direct stacking of Ti3C2 MXene nanosheets could outperform existing recording or stimulation electrodes in electrical properties.

In this article, we successfully fabricated recording and stimulation electrode fibers directly from stacked Ti3C2 MXene nanosheets for the first time. Within the electrode fibers and at the electrode-solution interface, the charge transfer rate is exceptionally high, reflecting high conductivity and low impedance. Doping the electrode fibers with PEDOT has further enhanced their electrical properties, resulting in superior recording and stimulation capabilities as demonstrated in electromyography (EMG), electrocardiograph (ECG), cortical recording and subthalamic nuclei deep brain stimulation (STN-DBS) experiments. Moreover, the manufacturing process of these electrode fibers is straightforward and cost-effective.

## Fabrication process

The fabrication process can be divided into three main parts: ink preparation, wet spinning, and solidification. During the ink preparation process, 1 ml of Ti3C2 MXene aqueous dispersion (5 mg/ml) is mixed with 100 µl of PEDOT-PSS aqueous dispersion (11 mg/ml). The mixture was shaken vigorously and then further dispersed using ultrasonication. Subsequently, the ink was loaded into a syringe and extruded through a pneumatic dispensing machine into a rotating tray containing an acetic acid-chitosan coagulation bath. The shear force applied by the turntable induced the alignment of the ordered liquid crystal MXene domains (Fig. 1A). This aligned state was retained to form a coarse and water-rich filament during the coagulation processes due to the slow absorption of water in the ink by chitosan and acetic acid (Fig. 1B). Following the wet spinning process, these coarse filaments were removed from the coagulation bath using tweezers, washed in a dish containing pure water. Then, the filaments were laid out on a drying rack for solidification at room temperature, forming flexible fibers with a diameter of approximately 30 µm, named MPP (Fig. 1C). The fibers are stored in a vacuum container for preservation and further drying. To study the effects of PEDOT-PSS, similar fibers are prepared using a 5 mg/ml Ti3C2 MXene aqueous dispersion following the same procedure, named MX.

**Fig. 1.**
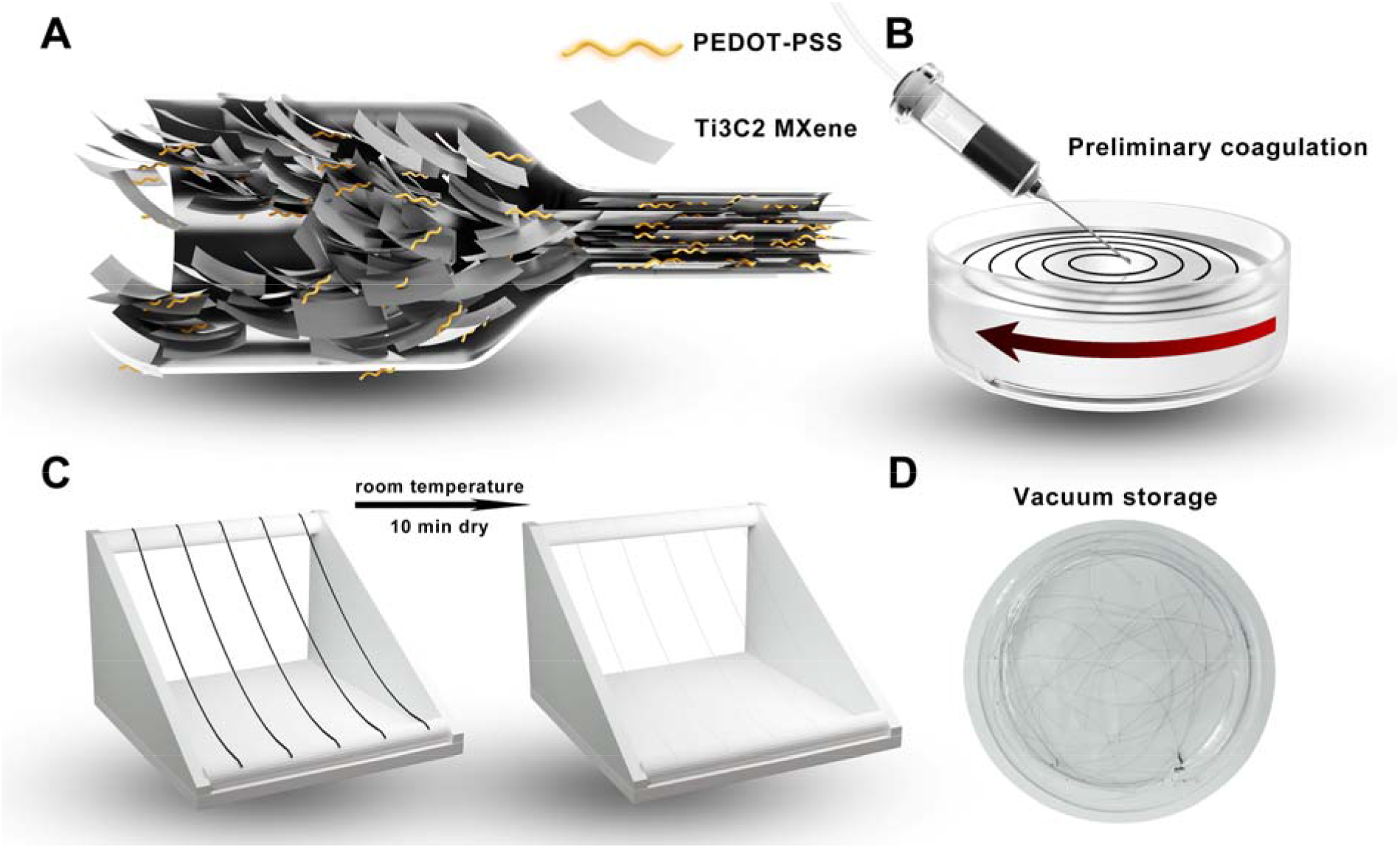
Fabrication process. **(A)** The alignment of MXene sheets under the shear force in the needle. **(B)** Setup of the coagulation process. **(C)** Schematic representation of the drying process. **(D)** The fiber electrode collected in a dish.

## Materials chatacterization

The morphology of MX and MPP fibers was examined using scanning electron microscopy (SEM) (Fig. 2A and 2C). The MPP fibers feature radial wrinkles along their sides and exhibit a complex, porous structure at the cross-section, which is similar to that of MX fibers. SEM analysis of the fiber cross-section revealed that the MXene sheets are oriented along the fiber direction, which may enhance the electrical properties of the fibers[16, 17]. The surface morphology of these fibers is similar to that of MXene-based fibers previously studied in supercapacitors [18, 19]. The similarity suggests that the wet spinning process effectively imparts a structured, porous texture to fibers based on layered materials, which may lead to excellent capacitive properties. It is noteworthy that the fibers prepared in this study exhibit a more finely ordered arrangement of nanosheets, which is attributed to the contributions from gradual fractional solidification [19]. In addition, the manual factors during the drying process can result in fibers with varying cross-sectional shapes (Fig. 2A right and 2C right). Furthermore, the distribution of MXene sheets and the PEDOT-PSS molecular chains appear to be homogeneous in MPP fibers, as demonstrated in the energy dispersive spectroscopy (EDS) map for titanium (Ti) and sulfur (S) (Fig. 2B and 2D). The EDS spectrum and semi-quantitative analysis confirmed the presence of S in the MPP, further verifying the doping of PEDOT-PSS (Fig. S1). The fibers exhibited excellent flexibility, easily wrapping around a glass rod with a diameter of 6 mm (Fig. 2E).

**Fig. 2.**
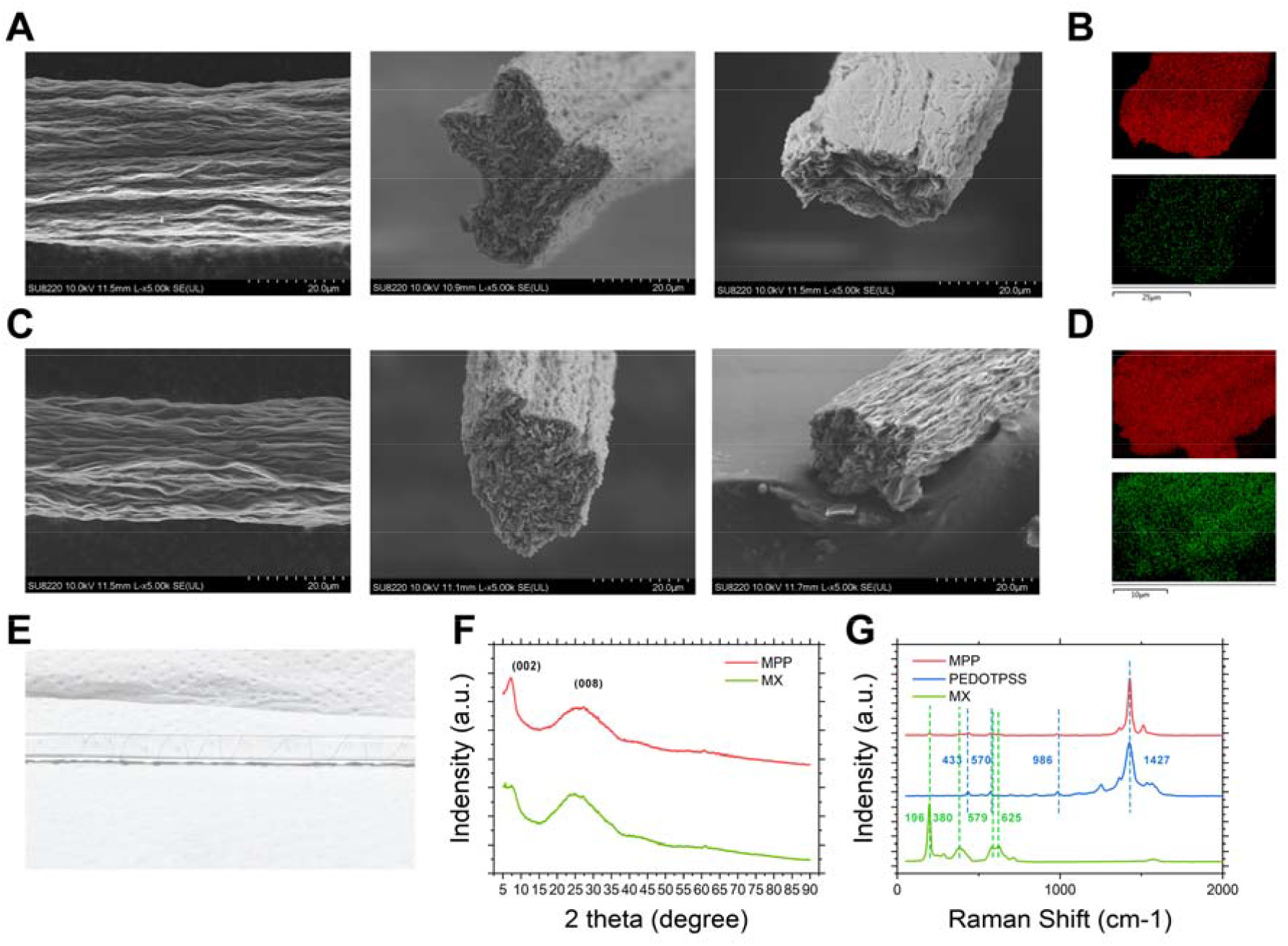
Materials characterization. **(A)** SEM micrographs of MX fibers. Side(left), ideal section(middle), example of deformation (right). **(B)** EDS mapping for Ti and S of MX fiber. **(C)** SEM micrographs of MPP fibers. **(D)** EDS mapping for Ti and S of MPP fiber. **(E)** MPP fiber wrapping around a glass rod with a diameter of 6 mm. **(F)** XRD patterns of MX and MPP. **(G)** Raman spectra of MPP, PEDOT-PSS and MX.

In the X-ray diffraction (XRD) result, similar diffraction patterns were observed for both MX and MPP samples (Fig. 2F). A sharp (002) peak at 7.36 ° and a broader (008) peak at 25.92° were identified which are similar to previous research [20, 21]. The two board peaks indicate two different spacings of nanosheet arrangements at 12.00 A and 3.43 A. Upon the addition of PEDOT-PSS, an increase in peak intensity at the (002) plane was observed in the MPP samples, while the intensity of the (008) peak slightly decreased. This indicates that the incorporation of PEDOT-PSS facilitates a partial transition of the tightly packed nanosheets from the (008) arrangement to the (002) arrangement. It suggests that PEDOT-PSS lines are dispersed among the MXene layers, forming new conductive pathways.

Raman spectroscopy was employed to assess the doping effects of PEDOT-PSS on MXene (Fig. 2G). The characteristic peaks of MX fibers are observed between 200-800 cm-1, consistent with existing literature [11, 22, 23]. PEDOT-PSS exhibits bands at 986 and 436 cm-1, attributed to the deformation of the C–O–C bond and the oxyethylene ring, respectively. The hybrid MPP fiber displayed combined characteristics of both MX and PEDOT-PSS spectra. Notably, the peak at 1427 cm-1, associated with the degree of PEDOT-PSS doping, remained constant in the MPP fibers, indicating that the presence of MXene did not significantly alter the doping level of PEDOT-PSS. In the composite fibers, the peak at 1427 cm-1 was pronounced, signifying the presence of PEDOT-PSS. The corresponding peaks for MXene became extremely weak, likely due to the uniform dispersion of PEDOT-PSS within the composite fibers, which absorb the scattered waves from MXene. This phenomenon has also been observed in a previous study [23], albeit to a lesser extent. This difference may be attributed to a more uniform distribution of PEDOT within the MPP fibers compared to the earlier study.

The linear voltammetry results obtained from 1 cm long electrode fibers were used to calculate the electrical conductivity (Fig. 3A). The conductivity of MX is 9.32 ± 1.32 × 10^4^ S/m (mean ± SD, n = 6), and the conductivity of MPP is 2.16 ± 1.46 × 10^5^ S/m (mean ± SD, n = 6). Both materials demonstrate outstanding electrical conductivity, which can reduce signal loss during transmission between electrodes due to their very low volumetric resistivity. To facilitate the evaluation of charge transfer properties and impedance, the fibers were encapsulated using silica tubing and light-curable dental cement, exposing approximately 300 µm (∼28981 µm^2^) (Fig. S2). The encapsulated samples were then subjected to electrochemical measurements in 0.1 M phosphate buffered saline (PBS). Specifically, we conducted Electrochemical Impedance Spectroscopy (EIS), Cyclic Voltammetry (CV), and voltage-transient experiments to measure impedance, CSCc, water window, and CIC.

**Fig. 3.**
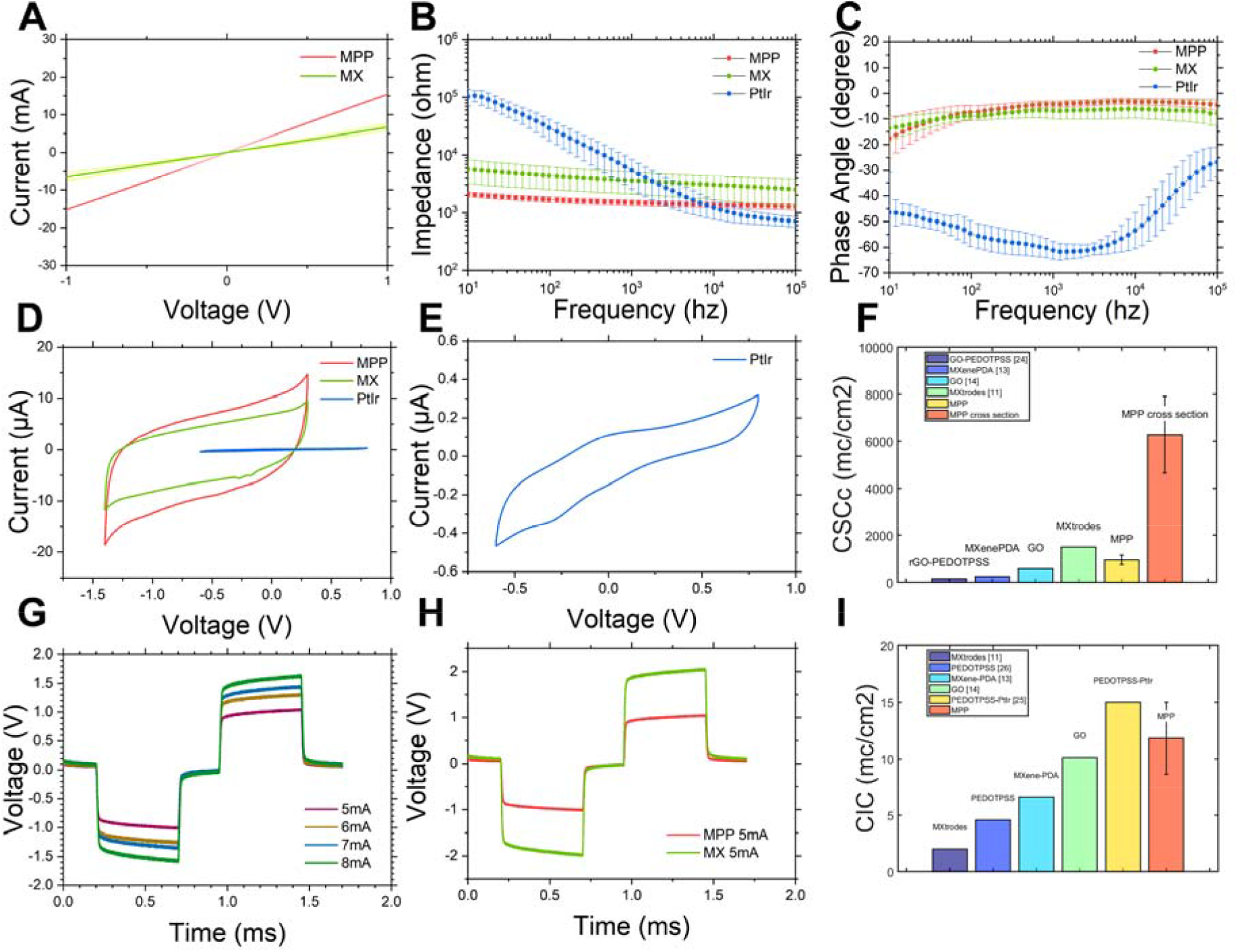
Electrical properties. **(A)** linear voltammetry results obtained from 1 cm long electrode fibers. **(B)** Impedance modulus of MPP, MX and PtIr samples. **(C)** Phase angle of MPP, MX and PtIr samples. **(D)** CV curves (50mv/s) of MPP, MX and PtIr samples. **(E)** Detail of the PtIr CV figure. **(F)** CSCc of different electrode materials. “MXtrodes” is made of Ti3C2 Mxene and cellulose cloth. **(G)** Voltage transients in response to biphasic current pulses, with tc = ta = 500 μs and tip = 250 μs, current amplitudes ranging from 5 mA to 8 mA. **(H)** Voltage transients of MPP and MX under 5 mA current. **(I)** CIC of different electrode materials.

The results of EIS revealed that the MXene-based fibers exhibited significantly lower impedance compared to the PtIr electrodes at frequencies below 1 kHz (Fig. 3B). At 10 Hz, the impedance values of the MPP fibers are 2068.50 ± 443.77 (mean ± SD, n = 6), which are 36% of the impedance of MXene electrodes (5761.67 ± 1053.25, mean ± SD, n = 6) and approximately 50 times lower than that of the PtIr electrodes (103663.33 ± 32793.19, mean ± SD, n = 6).

Compared to the PtIr electrodes, the MXene-based electrodes exhibit a larger phase angle (Fig. 3C), indicating a reduced imaginary component of impedance, possibly due to a larger electrode surface area and greater double-layer capacitance. Furthermore, compared to MXene electrodes, the MPP electrodes demonstrate a larger phase angle at frequencies above 40 Hz, which may be attributed to the additional capacitance provided by the PEDOT.

In CV tests at 50mv/s (Fig. 3D and 3E), the electrochemical stability window for MPP fibers determined from wide-scan CVs is -1.4v-0.3v, which is similar to previous research about MXene based electrodes [11, 13]. This wide potential range is advantageous for therapeutic electrical stimulation applications and the stimulation waveforms can be engineered to minimize voltage excursions in the anodic range while taking advantage of the large cathodic limit. The CV curves of the MXene-based electrode exhibited a nearly rectangular shape with no redox peaks observed which suggested that the electrochemical interaction at the electrode-electrolyte interfaces is controlled by capacitive rather than Faradic process. Additionally, compared to MX electrodes, MPP electrodes displayed enhanced capacitive behavior, which is advantageous for recording electrophysiological signals. The CSCc of the MPP fibers is 989.77 ± 174.03 mC/cm^2^ (mean ± SD, n = 6), which is substantially larger than that of MX fibers (626.24 ± 126.61 mC/cm^2^, mean ± SD, n = 6) and approximately 100 times greater than that of PtIr electrodes (9.18 ± 2.95 mC/cm^2^, mean ± SD, n = 6). Notably, the CSCc of the cross-sectional area of MPP fibers is 6278.99 ± 1634.22 mC/cm^2^ (mean ± SD, n = 6), significantly higher than the CSCc obtained from the 300 µm samples tested. Compared to the crumpled lateral surfaces, the cross-section features densely packed nanoplatelets forming a porous structure, greatly enhancing the actual contact area with the electrolyte, which is considered the reason for the significant increase in CSCc. Compared to recent research findings, the CSCc of MPP is outstanding (Fig. 3E) [11, 13, 14, 24].

Next, we evaluated the voltage transients generated on each electrode when delivering charge-balanced, cathodic-first biphasic current pulses. Each phase of the pulse had a duration of 500 µs (tc and ta), with an interpulse interval of 250 µs. The CIC of the MPP fibers was calculated based on the voltage transients with different currents (Fig. 3G). When the current is 5 mA, MPP revealed a lower voltage compared to MX (Fig. 3H). The CIC of MPP fibers measured at 11.81 ± 3.55 mC/cm^2^ (mean ± SD, n = 5), which is higher than that of MX fibers (5.90 ± 1.67 mC/cm^2^, mean ± SD, n = 5) and approximately 200 times greater than that of PtIr electrodes (0.06 ± 0.01 mC/cm^2^, mean ± SD, n = 5). Compared to recent stimulation electrode materials, the CIC of MPP is also at the forefront (Fig. 3I) [11, 13, 14, 25, 26].

It is believed that the more ordered alignment of MXene nanosheets due to the wet spinning process improves the efficiency of electron transport between the nanosheets. Additionally, PEDOTPSS intercalated between MXene layers provides extra conductive pathways, further enhancing electron transport efficiency and augmenting capacitive behavior in the solution system.

To further evaluate MPP electrodes, we investigated their electrochemical stability. First, conventional STN-DBS stimulation (130 Hz, t_cathodic_ = t_anodic_ = 100 μs, I= 90 μA) was applied to the MPP electrodes in the PBS solution system. The voltage curves of the electrode were examined after stimulation number is 1, 10k, and 100k (Fig. 4A). Additionally, we conducted 1000 CV tests on the MPP samples (Fig. 4B). The MPP samples maintained a high capacitance retention of 93.33 ± 3.20% (mean ± SD, n = 4) after 1000 CV cycles. The electrodes did not show abnormal voltage behavior after 100 k stimulation and demonstrated good stability in the 1000 CV. The excellent electrochemical stability might be attributed to the lower current densities at the electrode surface, akin to previously reported electrodes with rough surface topographies. This type of stability is crucial for applications where electrodes are subjected to repeated electrical loading [11].

**Fig. 4.**
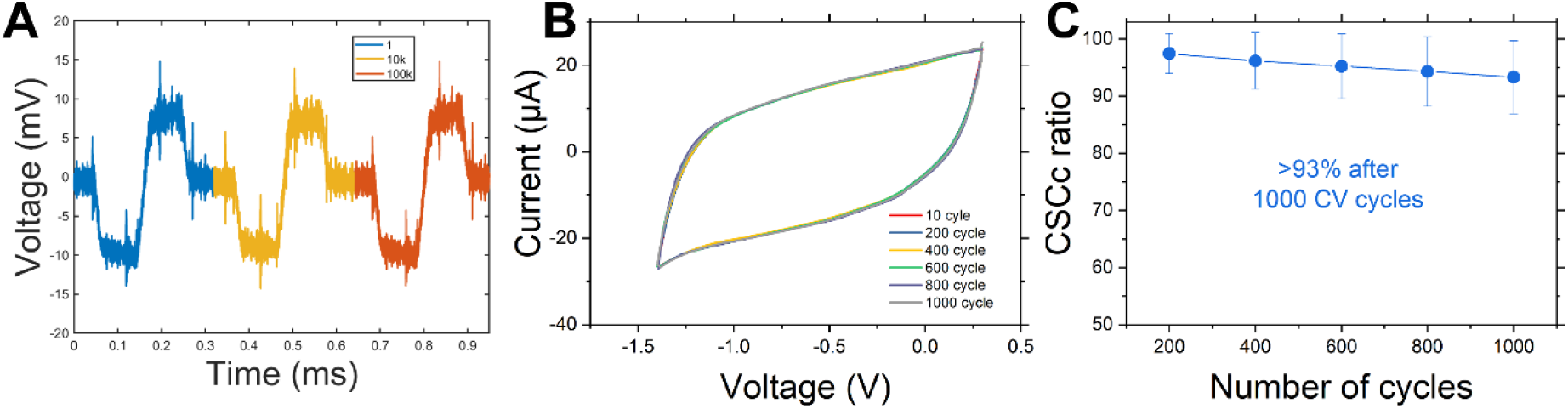
Electrochemical stability. **(A)** Voltage transients in response to biphasic current pulses, with t_cathodic_ = t_anodic_ = 100 μs, I= 90 μA, the stimulation numbers are 1, 10k and 100k. **(B)** 1000 cycles CV curves (100mv/s) of MPP. **(C)** CSCc retention after different recycling CV scanning.

The safety and compatibility of MXenes with neuronal cells have been widely validated in numerous neural interface studies recently [12, 27]. Based on the previous research, Cytotoxicity tests were conducted on primary cortical neurons using a 5mm segment of MPP electrodes in a 96-well plate. Optical images demonstrated healthy cell growth, with a minor adhesion to the MPP after 24 hours and significant adhesion after 72 hours (Fig. S3A). The viability assessments at 24 and 72 hours revealed no significant differences in cell survival between Ti3C2 and control blank cultures, with cell survival rates of 108.07 ± 8.46% (mean ± SD, n = 6) after 24 hours and 110.97 ± 18.88% (mean ± SD, n = 6) after 72 hours, indicating biocompatibility (Fig. S3B).

## Recording test on human skin surface

To further assess the recording capabilities of the MPP, an MPP fiber was affixed to a commercial Ag/AgCl electrode, which had its gel layer removed, using Polyimide (PI) tape, exposing only 5 mm of the fiber as the recording area (Fig. 5a). The measurements of skin impedance were carried out by adhering a pair of electrodes on forearm skin with a distance of 2 cm (Fig. 5B). Even with a significant difference in the solid electrode area (MPP= 0.0047 cm2, commercial electrode= 3.14 cm2), the MPP electrode demonstrates lower impedance when the frequency below 100 Hz (Fig. 5C).

**Fig. 5.**
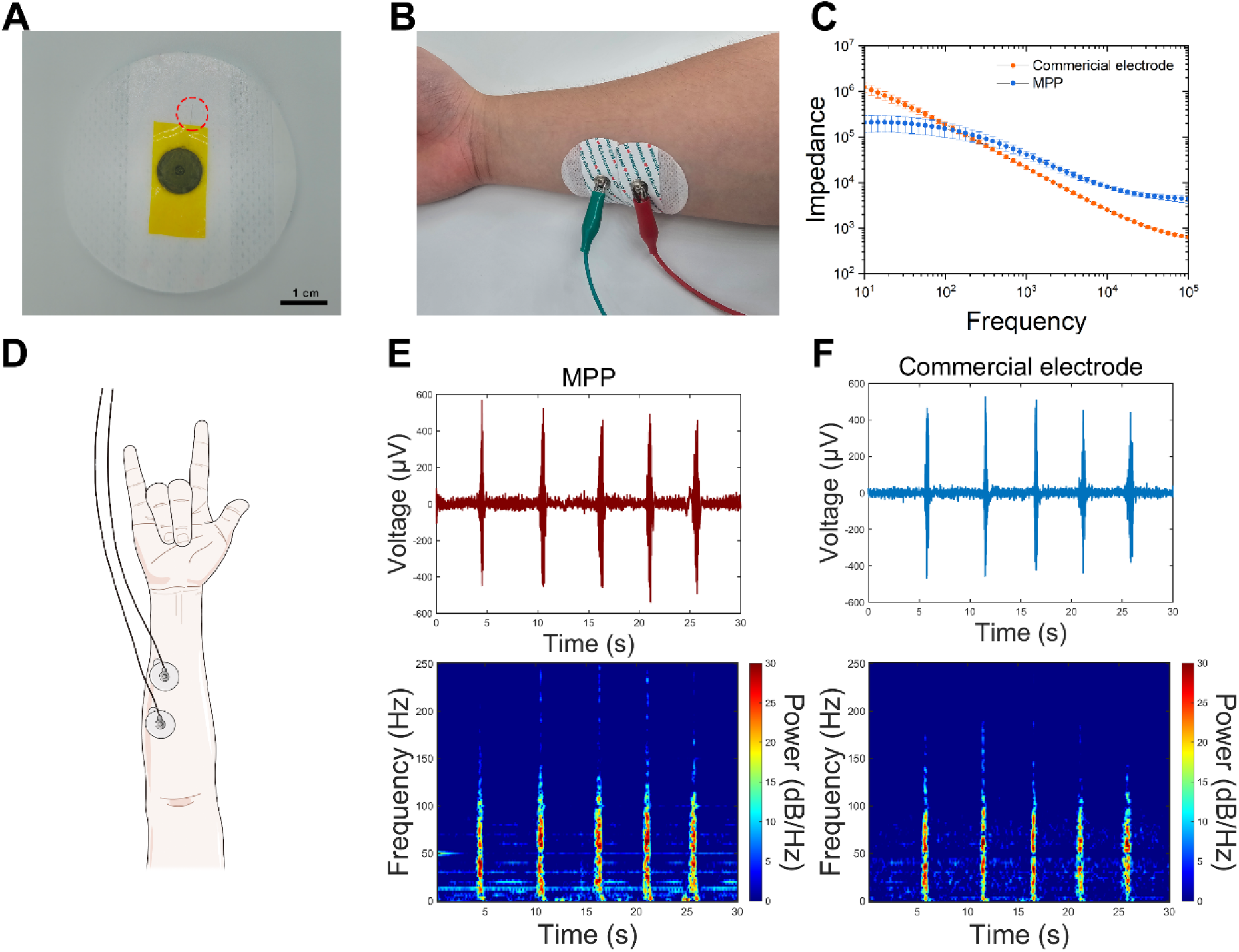
EMG detection on brachioradialis. **(A)** Optical images of MPP EMG electrode. **(B)** Set up of skin impedance test. **(C)** Skin impedance modulus of commercial electrodes and MPP samples. **(D)** Illustration of the specific gesture and setup. **(E)** EMG recordings and spectrogram of MPP. **(F)** EMG recordings and spectrogram of commercial electrode.

**Fig. 6.**
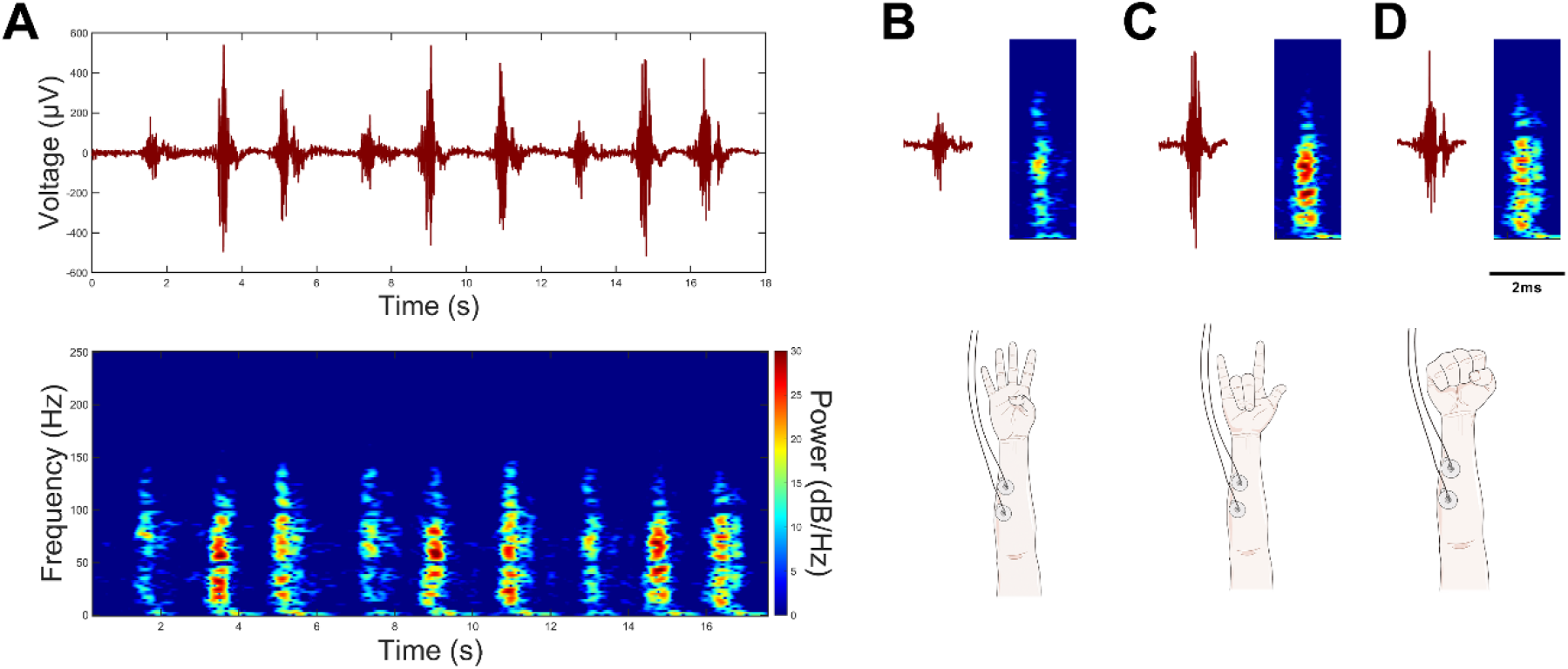
Recordings of different gestures. **(A)** Recordings and spectrogram of three gestures repeated continuously. **(B)** Detailed recordings and spectrogram of gesture 1. **(C)** Detailed recordings and spectrogram of gesture 2. **(D)** Detailed recordings and spectrogram of gesture 3.

The brachioradialis muscle is one of the main elbow flexors along with the biceps brachii and the brachialis. In neurological patients, brachioradialis is often affected by structure and muscle activity alterations, leading to pathological upper limb patterns of movement, and therefore is one of the main targets of detection[28]. According to previous studies[29], three identical electrodes were used: one placed on the back of the hand as a reference electrode, and the other two placed on the brachioradialis as the working electrode group (Fig. 5D). Signals were elicited by specific gestures, with each elicitation spaced more than 5 seconds. In the waveform graph, the evocation of the EMG signal can be observed, and its shape is similar to that found in a previous study[29]. The SNR of the MPP electrode is 25.78 ± 1.47 dB (mean ± SD, n = 4), which is similar to the SNR of the commercial electrode (26.74 ± 1.32 dB, mean ± SD, n = 4).

Additionally, in the frequency spectra of both types of electrodes, it is clear that upon gesture elicitation, the EMG signals lasted about one second, with frequencies mainly distributed between 10 to 140 Hz.

We further investigated the EMG signal variations from the brachioradialis muscle induced by different gestures using the MPP electrode. It was observed that the EMG signal elicited by the first gesture was relatively weak, with the signal frequency predominantly ranging between 50-100 Hz. The second gesture notably stimulated the brachioradialis, producing a strong EMG signal across the frequency range of 20-100 Hz. The EMG signal strength induced by the fist-clenching gesture was greater than that of the first gesture but less than the second. The frequency distribution was similar to that of the second gesture, albeit with slightly lower intensity and longer duration. In the waveform plot, the fist-clenching gesture was characterized by a negative peak following the main peak, likely due to the extensive muscle resetting during the release of the gesture.

The activity of the sympathetic nervous system is closely related to arousal and stress, and it aids in understanding the mechanisms within human cardiovascular physiology and pathophysiology[30]. We also tried to detect surface electrophysiological signals induced by sympathetic excitation of the hand. The recording electrode was placed at the thenar eminence of the non-dominant hand, the reference electrode was placed on a less obtrusive location, at the ventromedial forearm approximately 15 cm below the hand electrode, the grounding electrode was placed at the dorsum of the hand [31]. After the volunteers entered a resting state, an alarm clock set to be excited approximately two minutes later was placed beside the head, and then the recording started. In the typical recording result, we can see that before the arousal, the thenar eminence had only occasional weak signals, while after the arousal, there was a continuous burst of electrical signals recorded by the MPP electrode. The signals concentrated in the range of 0-10hz, and a small number of high-frequency signals appeared in the range of 0-80hz (Fig. S4).

ECG signals in the V2 area were detected by an MPP electrode and a commercial electrode (Fig. 7A). Both electrodes captured nearly identical QRS waveforms, consisting of a small positive wave (r), followed by a large negative wave (S), and a large positive wave (R’). A faint P wave can be observed approximately 0.12 ms before the QRS waveforms, and a symmetric T wave inversion is observed approximately 0.15 ms after the QRS waveforms. (Fig. 7B). In the spectrogram, the duration of each excited signal was ∼0.6 ms, composed of a ∼0.4 s high-frequency signal followed by a ∼0.2 s low-frequency signal (Fig. 7C and 9D). The results of the two electrodes were also similar. Notably, compared to the commercial electrode, the MPP electrode displayed additional high-frequency signals in the 50-70 Hz range, which may be due to its small recording sites.

**Fig. 7.**
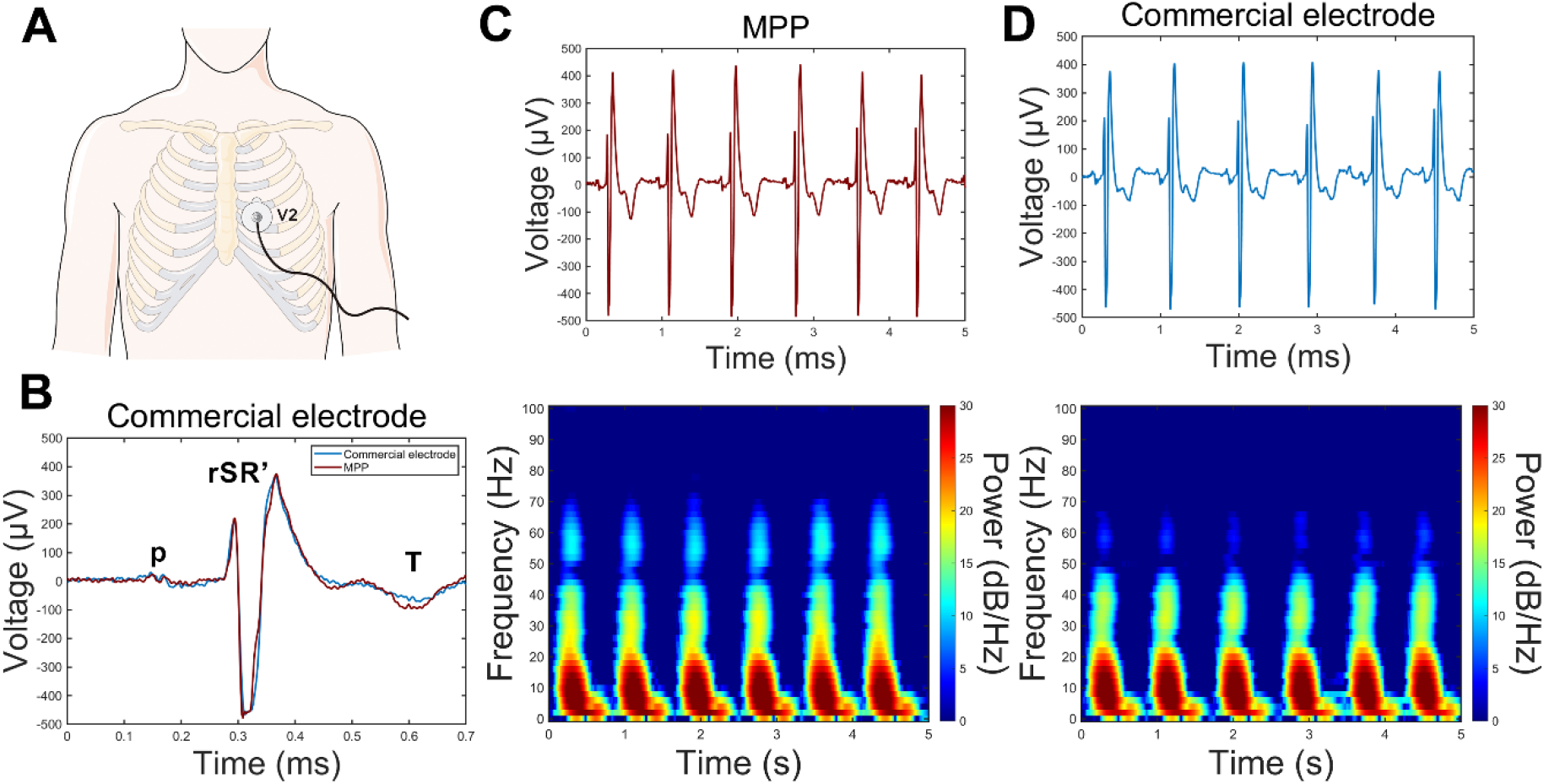
ECG detection on V2. **(A)** Illustration of the ECG detection setup. **(B)** ECG waveforms were recorded on the two electrode types, marked with salient ECG features. **(C)** Five seconds of ECG recordings and spectrum of MPP electrode. **(D)** Five seconds of ECG recordings and spectrum of the commercial electrode.

## In vivo recording of the primary visual cortex

To evaluate the in vivo electrophysiological recording capabilities of the electrode fiber, two encapsulated MPP electrodes were assembled in parallel with a laterally insulated PtIr fiber (diameter = 35 µm) to form a recording electrode group. This electrode group was implanted into the left primary visual cortex (V1, 2.7 mm posterior to bregma, 2.4 mm lateral to the midline) of 9-week-old C57BL/6 mice (weighing 22-28 grams). Visual stimulation was presented to the right eye of the subjects using a computer monitor (Fig. 8A). Each stimulation consisted of a 3-second contrast-reversing checkerboard pattern at 1 Hz, interspersed with a 12-second gray background. In typical recording results, noticeable changes in the waveform in the V1 area were observed during screen stimulation (Fig. 8B). The results from the Peristimulus Time Histogram (PSTH) and spectrogram also demonstrated signal bursts during stimulation (Fig. 8C). During the stimulation periods, peaks significantly exceeded the average, with signal power markedly increasing and frequency distribution shifting from 0-10 Hz to 0-40 Hz. During non-stimulated periods, only sporadic spike signals and low-frequency faint signals were observed. The recording results from the MPP electrodes are similar to those from the PtIr, displaying a stronger signal power in the frequency spectrum, indicating excellent recording performance.

**Fig. 8.**
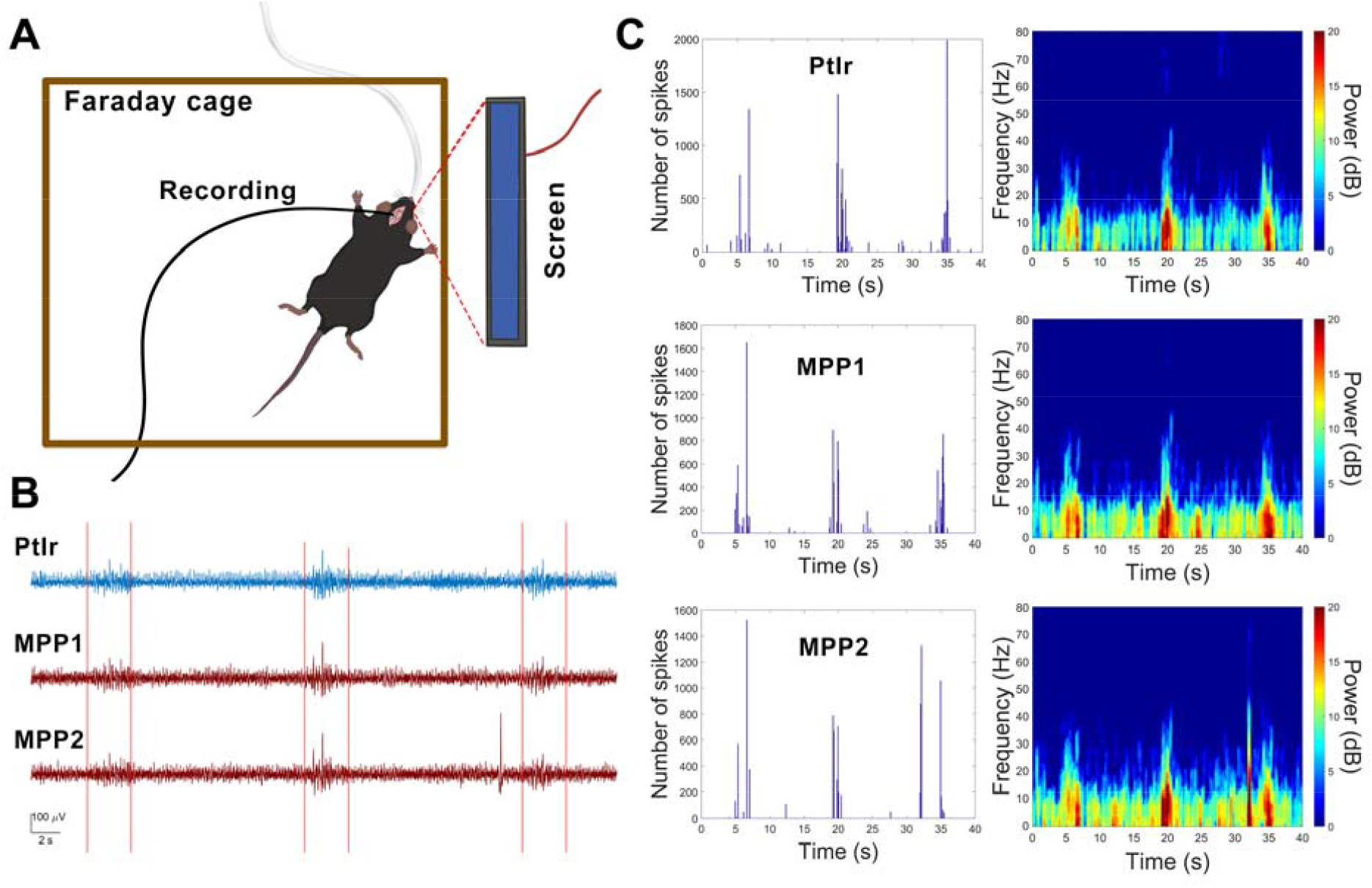
Example resting state and evoked recording from one animal. **(A)** Illustration of the setup. **(B)** Example channel of the PtIr and MPP. The red line indicates the stimulation periods. **(C)** PSTH and spectrogram of PtIr and MPP electrodes.

## In vivo STN-DBS and MRI imaging

STN-DBS is a surgical treatment commonly used for managing symptoms of Parkinson’s disease. The stimulation capacity of MPP microelectrodes was demonstrated through STN-DBS in Sprague-Dawley (SD) rats (Fig. 9A). The bipolar stimulation electrode assembly consisted of platinum-iridium and MPP wires, with the ends of the wires exposed by 300 µm to serve as the stimulation sites. Magnetic resonance imaging (MRI) was used to confirm the placement of the electrodes (Fig. 9B). Based on the latest rat three-dimensional brain atlas database [32], we used the QuickNII tool to register sections near the electrode implantation site within the MRI three-dimensional scan data [33, 34]. After comparison with the standard atlas, the electrode’s terminal was precisely implanted at the STN location (Fig. 9C). The results indicated that the MPP electrodes are clearly in MRI scans, which can be attributed to the magnetic susceptibility properties of MXene. The magnetic susceptibility properties are similar to those of human tissue, enhancing their compatibility and visibility in clinical imaging applications [35].

**Fig. 9.**
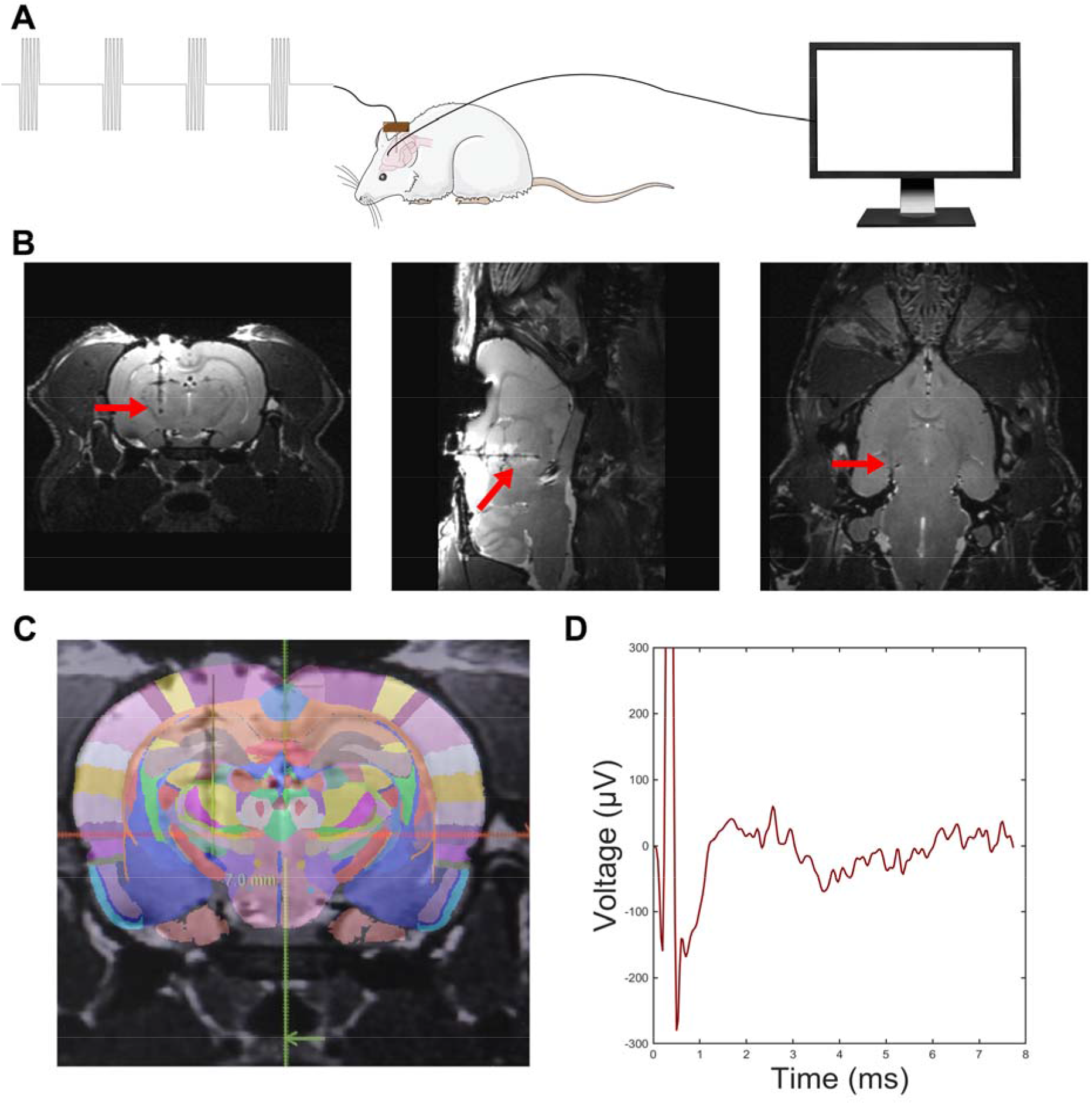
STN-DBS test. **(A)** Illustration of the setup for the STN-DBS on SD rat. **(B)** Representative coronal (left), sagittal (middle) and horizontal (right) sections of the T2 MRI images of rat brains implanted with an electrode group. The red arrow points to the electrode. **(C)** Schematic illustration following localization based on the 3D brain atlas. **(D)** Average evoked LFP after 800 stimulations.

After confirming the implantation sites, high-frequency stimulation consisting of 130 Hz square constant current pulses (biphasic, symmetric, charge-balanced pulses at 90 μA and 100 μs width per phase) was applied to the MPP-PtIr bipolar microelectrodes. Each stimulation session consisted of 200 pulses, manually triggered, with intervals exceeding five seconds between sessions. The evoked potential signals were recorded through titanium screws implanted in the skull near the motor cortex. Noise unrelated to the electrical stimulation was eliminated by averaging 800 segments of 8 ms evoked potential, resulting in the identification of characteristic peaks following the stimulation (Fig. 5d). We observed response voltages of approximately 50 µV at ∼1.67 ms and 2.57 ms after the stimulation, which are similar to previous researches [36-38].

## Discussion

As anticipated, electrode fibers constructed through the direct stacking of MXene nanosheets possess electrical performance that significantly surpasses that of other MXene-modified electrodes. Doping with PEDOT-PSS enhances the conductive pathways while preserving the original folded structure of the fibers, further improving their electrical properties. The CSCc of the MPP electrodes is 989.77 ± 174.03 mC/cm^2^, with the cross-sectional CSCc nearly seven times this value. Their exceptional recording potential has been validated in skin tests: MPP electrodes demonstrate recording capabilities nearly equivalent to commercial electrodes, despite having a nearly 1000-fold smaller surface area. Additionally, MPP exhibits a stronger signal-to-noise ratio than platinum-iridium wires in cortical recording experiments.

Their high CIC of 11.81 ± 3.55 mC/cm^2^ also highlights their stimulation capabilities, which have been confirmed in rat STN-DBS experiments. The MPP fibers, with their robust recording and stimulation abilities and MRI compatibility, hold broad prospects in the medical device field, particularly for invasive electrical neural interfaces. In the future, through advancements in the spinning process and exploration of array technologies, these directly stacked MXene electrodes are expected to further reduce the size of electrical interfaces and enhance the precision of recording and stimulation.

## Acknowledgments

ChatGPT-4 is used to assist in writing MATLAB scripts and editing English grammar.

## Funding

Scientific and Technological Innovation 2030 Key Project (2022ZD0209800) Natural Science Foundation of China Grants (31930047)

National Key R&D Program of China (2020YFC2008503) the Strategic Priority Research Program of Chinese Academy of Science (XDB32030103)

National Special Support Grant (W02020453)

NSFC-Guangdong Joint Fund (U20A6005)

Key-Area Research and Development Program of Guangdong Province (2018B030331001 and 2018B030338001)

Shenzhen Infrastructure for Brain Analysis and Modeling (ZDKJ20190204002)

## Author contributions

Conceptualization: SG

Methodology: SG, ZD, PL, SY

Investigation: SG, ZD, PL

Visualization: SG, SY

Funding acquisition: ZD, GB

Project administration: SG, ZD

Supervision: ZD, GB

Writing – original draft: SG

Writing – review & editing: SG, ZD

## Data and materials availability

All data are available in the main text or the supplementary materials.

