## Supplementary Materials for "Stacked MXene/PEDOT-PSS Electrode Fiber for High-Performance Recording and Stimulation"

Materials and Methods

Ti_3_C_2_ MXene was synthesized by selectively etching Al atomic layers from Ti_3_AlC_2_ MAX phase particles (<25 μm in size, XFNANO). The etching solution was prepared by dissolving 3.2 g of lithium fluoride (LiF, 99%, Sigma-Aldrich) in 40 mL of 9 M hydrochloric acid (HCl, Sigma-Aldrich) and stirring for 15 minutes. Subsequently, 2 g of Ti3AlC2 powder was gradually added to the etchant, and the reaction was allowed to continue for 48 hours at 40°C. The acidic dispersion was then repeatedly washed with deionized (DI) water through centrifugation (LuMickey-CHT210R) at 3500 rpm for 5 minutes per cycle until self-delamination occurred (pH ∼ 6). Following this, the dispersion was centrifuged at 3500 rpm for 30 minutes to separate the delaminated flakes from the multilayer MXene and unreacted MAX phase. The supernatant was then centrifuged at 5000 rpm for 1 hour, and the sediment containing the desired Ti3C2 flakes was collected. This sediment was freeze-dried for 24 hours to form a solid and then prepared into a 5 mg/mL dispersion.

In ink preparation process, 1 ml of MXene aqueous dispersion (5 mg/ml) is mixed with 100 µl of PEDOT-PSS aqueous dispersion (11 mg/ml, Shanghai Ouyi organic photoelectric Materials Co., LTD). The mixture is vigorously shaken and then sonicated for 5 minutes to further disperse it. Subsequently, the ink is loaded into a syringe equipped with a 27g needle, and a pneumatic dispensing machine supplies 5 kPa of air pressure to extrude the ink into an 8 rad/s rotating tray containing a coagulation bath. After a 10-minute coagulation period, the ink in the coagulation bath solidifies into coarse, water-rich filaments. Using flat-nosed plastic tweezers, one end of a filament is carefully grasped and extended by 10-20 cm before cutting off the end. This section is then gently placed into a dish filled with pure water to rinse off any residual coagulation fluid. Once washed, the filaments are spread out on a drying rack to air dry at room temperature, forming microfibers with an approximate diameter of 30 µm. These microfibers are subsequently further dried and stored in a vacuum container to preserve their quality and integrity.

The morphologies of the fiber electrode were examined by scanning electron microscopy (SEM) (QUANTA200, USA). To obtain structure information of fibers, the X-ray diffraction (XRD) patterns were measured with an XRD system (Smartlab, Rigaku, Japan) using Cu Kα radiation (λ = 1.54 Å) at a 2θ scan step of 0.02°. The Raman spectra were measured with a micro-Raman Spectrometer (LabRAM HR Evolutio, Horiba Jobin Yvon, Japan).

Before the electrochemical test, the tip of the fiber electrodes is adhered to using conductive silver glue and copper tape for easy connection with external circuits. The other end is insulated with silica tubing (inner diameter 100µm, outer diameter 170µm, TSP100170, MOLEX) and light-curable dental cement, leaving 300 µm exposed. The electrochemical properties of the fiber electrodes were studied using a three-electrode setup in 0.1 M PBS electrolyte, in which a single fiber was used as the working electrode, Ag/AgCl as reference electrode, and platinum mesh (1 cm × 1 cm) as counter electrode. CV and EIS were performed using the electrochemical workstation (CHI660E). Cyclic voltammetry was performed at a sweep rate of 50 mV/s. EIS measurements were carried out at an open circuit potential by applying an alternating-current voltage with an amplitude of 5 mV in a frequency range of 10 Hz to 100,000 Hz. The charge storage capacity (CSC) was calculated as the time integral of the cathodic current recorded over the potential range (MPP and MX: −1.4 - 0.3V, PtIr: −0.6-0.8V) in the second cycle

For voltage transient experiments, a two-electrode cell consist of electrode sample and platinum mesh was used. Cathodic-first biphasic current pulses with a duration of 500 μs in both phases and an interpulse interval of 250 μs were delivered to the tested sample with a stimulator (Flexon). Voltage transients under the current pulses were recorded with an oscilloscope, and the negative potential excursion (Vexc) was considered as the maximum negative voltage. The charge-injection-limit was calculated by multiplying the current amplitude and pulse duration at which Vexc reaches the water reduction limit (−1.4 and −0.6 V for MPP electrodes and PtIr electrodes, respectively), divided by the geometric surface area of the electrodes.

Stability testing under continuous overcurrent pulsing was performed by immersing the MPP electrodes in a cell filled with 1× PBS, pH 7.4 at room temperature. a two-electrode cell (the same as above) was used and the cell was sealed in order to avoid evaporation of the electrolyte. the stimulation protocol involved a cathodic-leading bipolar square wave with an amplitude of 90 μA and a phase duration of 100 μs, delivered at a frequency of 130 Hz. Voltage curves were recorded after 1, 10,000, and 100,000 stimulations. In the 1000-cycle CV test, the scan rate was set at 100 mV/s, with other settings consistent with standard cyclic voltammetry procedures.

In cytotoxicity test, a PC12 cell line was obtained from the American Type Culture Collection (ATCC). PC12 cells were maintained at 37 °C (5% CO2, 95% air) in RPMI-10 medium (Sigma) supplemented with 5% fetal bovine serum (Atlanta Biologicals), and 5% penicillin/streptomycin (Sigma). The fresh medium was replaced every other day. Confluent cells were harvested with 0.25% trypsin−EDTA solution (Invitrogen). After sterilization in a high-temperature and high-pressure autoclave, 5 mm length MPP electrodes are placed into individual wells of a 96-well plate. Then, PC12 cells are seeded onto the fibers at a density of 12,000 cells per cm². The PC12 cells proliferate without changing the culture medium. After 24 and 72 hours, cells were observed under a light microscope, and then the CCK-8 solution (Mei5 Biotechnology, Beijing, China) was added to each well. After CCK-8 solution cultured cells for 1 h, the absorbance was computed by operating a Microplate Reader (BioTek) at a wavelength of 450 nm. The results are analyzed by the Tukey test.

In recording test, the electrode impedance on the skin was measured at the electrochemical workstation (CHI660E), covering a frequency range from 10 Hz to 100,000 Hz with an applied voltage of 0.2 V. Measurements were conducted by adhering a pair of electrodes on the forearm skin, spaced 2 cm apart. To detect the surface electromyography (sEMG) signals, two working electrodes were placed on the brachioradialis, and one reference electrode was attached to the back of the hand. To minimize the influence of circuit factors on the signal-to-noise ratio (SNR), direct improvements were made on a commercial interface to assess the recording capability of MPP electrodes. A single electrode wire, using PI tape, was attached to an AG/AGCL commercial electrode stripped of its gel layer, exposing only 0.5 cm of the microfilament for recording. One electrode was placed on the back of the hand as the reference, and two others on the brachioradialis served as the working electrode group. SNR was measured by stimulating signals through specific gestures, with each stimulation separated by over 5 seconds. Gesture recognition was performed with about a second interval between each gesture type. For measuring bursts of electrical signals induced by sympathetic nerve excitation, the subject was in a quiet and comfortable setting, with the recording electrode placed on the thenar eminence of the non-dominant hand. The reference electrode was situated on the ventromedial forearm approximately 15 cm below the hand electrode, and a grounding electrode was placed on the back of the hand. An alarm set to trigger approximately two minutes later was placed next to the head, and electrophysiological signals were continuously recorded during this period. For ECG signal testing, the recording electrode was placed in the V2 area of the chest, the reference electrode on the left leg, and the grounding electrode connected to the right leg. All signals were recorded using commercial signal recording equipment (DueLite, sampling rate 512 Hz). The sEMG signals were digitally filtered in MATLAB using a fourth-order Butterworth filter (0.1–250 Hz), and ECG signals were similarly filtered (0.1–150 Hz). All signals were processed in MATLAB with notch filters to remove significant noise at 50 Hz, 100 Hz, and 150 Hz. SNRs were calculated using the equation: SNR (dB) = 20 log10(A_signal_/A_noise)_, where Asignal and Anoise represent the amplitude of the signal and background noise, respectively.

In V1 detection from visual stimulation, two encapsulated MPP electrodes were assembled in parallel with a laterally insulated PtIr fiber (diameter = 35 µm) to form a recording electrode group. The groups were connected to recording equipment (Intan, sampling rate 30,000 Hz) using silver paint and copper tape. A single electrode group was implanted in the left primary visual cortex (V1, 2.7 mm posterior to bregma, 2.4 mm lateral to the midline) of 9-week-old C57BL/6 mice (weighing 22-28 grams). Induction was achieved with 3% isoflurane in oxygen at a flow rate of 0.5 L/min, then maintained at 1.5%. Artificial tears were used to prevent corneal drying and damage. After positioning the animal in a stereotaxic frame equipped with ear bars, the skin and connective tissue on the skull surface were removed. A small pinhole craniotomy was performed over the visual cortex using a high-speed dental drill, and bone fragments were carefully removed with forceps. Saline was continuously applied to the skull to dissipate heat from the drill. The procedure was conducted within a Faraday cage to facilitate subsequent stimulation and recording operations. A computer monitor was placed outside the cage to present visual stimulation to the right eye of the subjects. Each stimulation consists of a 3s contrast-reversing checkerboard pattern at 1 Hz, interspersed with a 12s gray background. A photosensitive resistor kit was placed on the monitor to synchronize the timing of the stimuli. All signals were digitally filtered in MATLAB using a fourth-order Butterworth filter (1–300 Hz) to isolate local field potentials (LFP). The recorded evoked unit activity was visualized by post stimulus time histograms (PSTHs). These histograms were used to visualize the rate and timing of neuronal spike discharges to an external stimulus. Peaks above the threshold, mean ±3 times the standard deviation (SD), were marked as spikes.

10-week old male Sprague-Dawley rats (Charles River) weighing 300–350 g were utilized to evaluate the in vivo stimulation performance of MPP fiber. Animals were anesthetized with 3.0% isofluorane in 0.8 L/min oxygen for 5 min prior to surgery and then maintained for the duration of the procedure at 2.25% isofluorane. Anesthesia level was monitored closely during the procedure by observing changes in respiratory rate, heart rate, body temperature (37.7 °C) and absence of the pedal reflex. Animals were placed in a stereotaxic frame and the hair was removed over the incision site. The working electrode group consists of one MPP fiber and one PtIr fiber, each with 300 µm exposed after encapsulation. Further encapsulation with sucrose is employed to prevent deformation and damage to the electrodes. The electrode group was implanted into the STN of rat (3.8 mm posterior to bregma, 2.5 mm lateral to midline, 7 mm deep from brain surface) through a craniotomy to perform DBS and evoke reliable EEG LFP signals as outlined in previous studies. The dura mater was cut to prevent possible separation of the electrode group. An EEG recording screw was placed on the motor cortex (+2.5 mm anterior to bregma, +2.5 mm lateral to midline) and the EEG ground screw was positioned above the cerebellum (0.8 mm posterior to lambda and 2.0 mm lateral to midline). The setup is secured by dental cement, exposing only a little part for connection. MRI was performed in a 9.4 Tesla, 30 cm diameter bore magnet MRI scanner (uMR 9.4T, United imaging Life Science Instruments, China). Rats were anesthetized with 4% isoflurane. During MRI scanning, isoflurane (0.5%) delivered via a nose cone to maintain anesthesia. Based on the existing brain atlas database [32], we used the QuickNII tool to analyze MRI results and determine the implantation site of the electrodes [33, 34]. Cathodic leading bi-polar square-wave stimulation with 90 μA amplitude and 100 μs duration per phase was delivered at 130 Hz to evoke LFP in motor cortex. All signals obtained by a commercial signal recording equipment (Intan, sampleing rate 30,000 hz). A total of 800 waveforms were averaged to obtain the mean evoked LFP from the stimulation.

Supplementary Figure

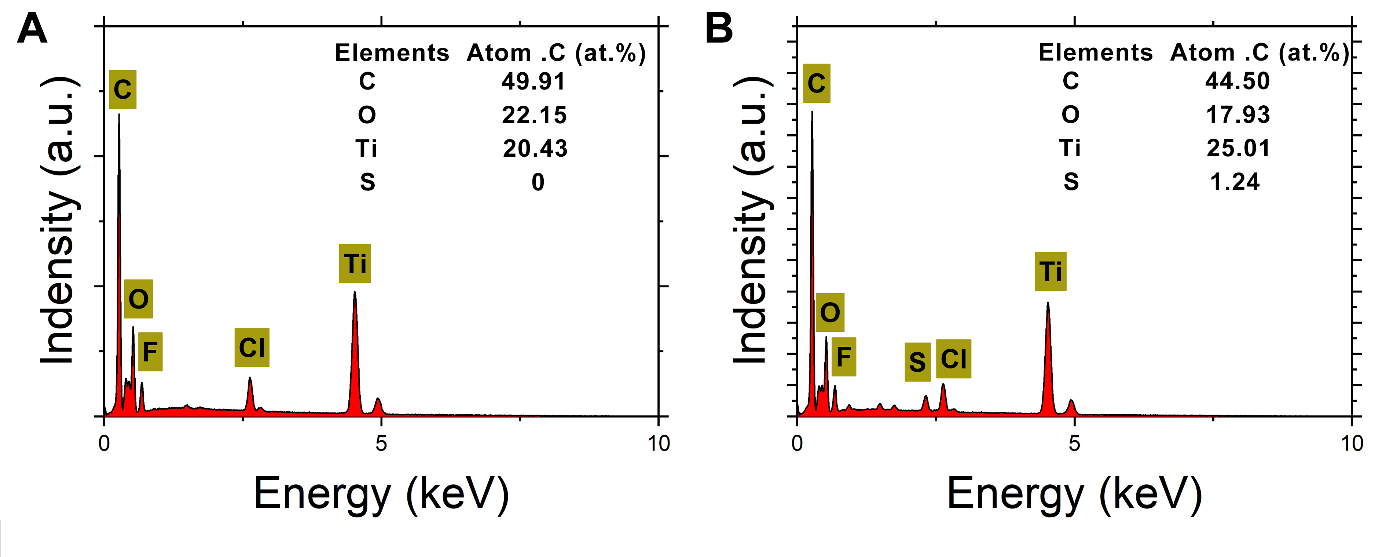

**Fig. S1.** EDS spectrum for **(A)** MX fibers. **(B)** MPP fibers.

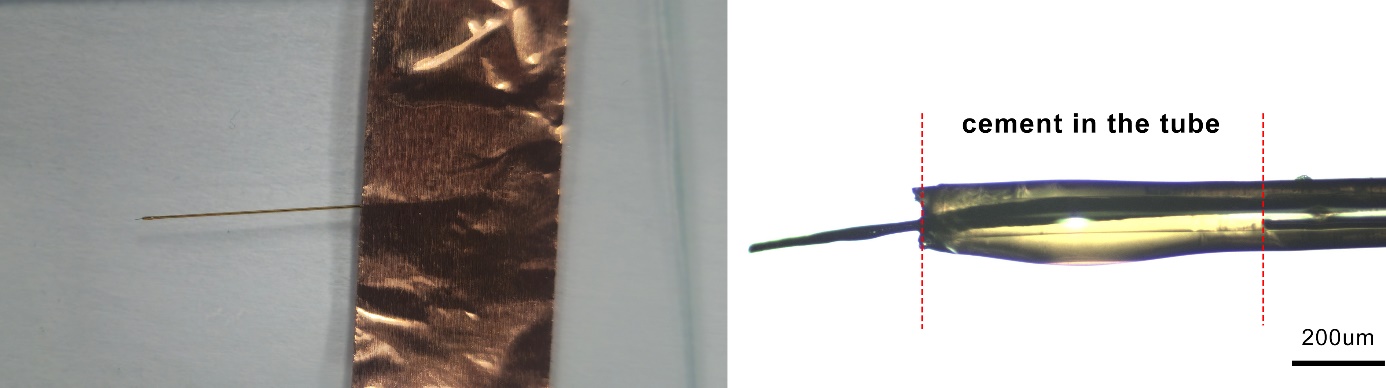

**Fig. S2. Optical images of packaged samples.** **(A)** Real product. **(B)** Detailed.

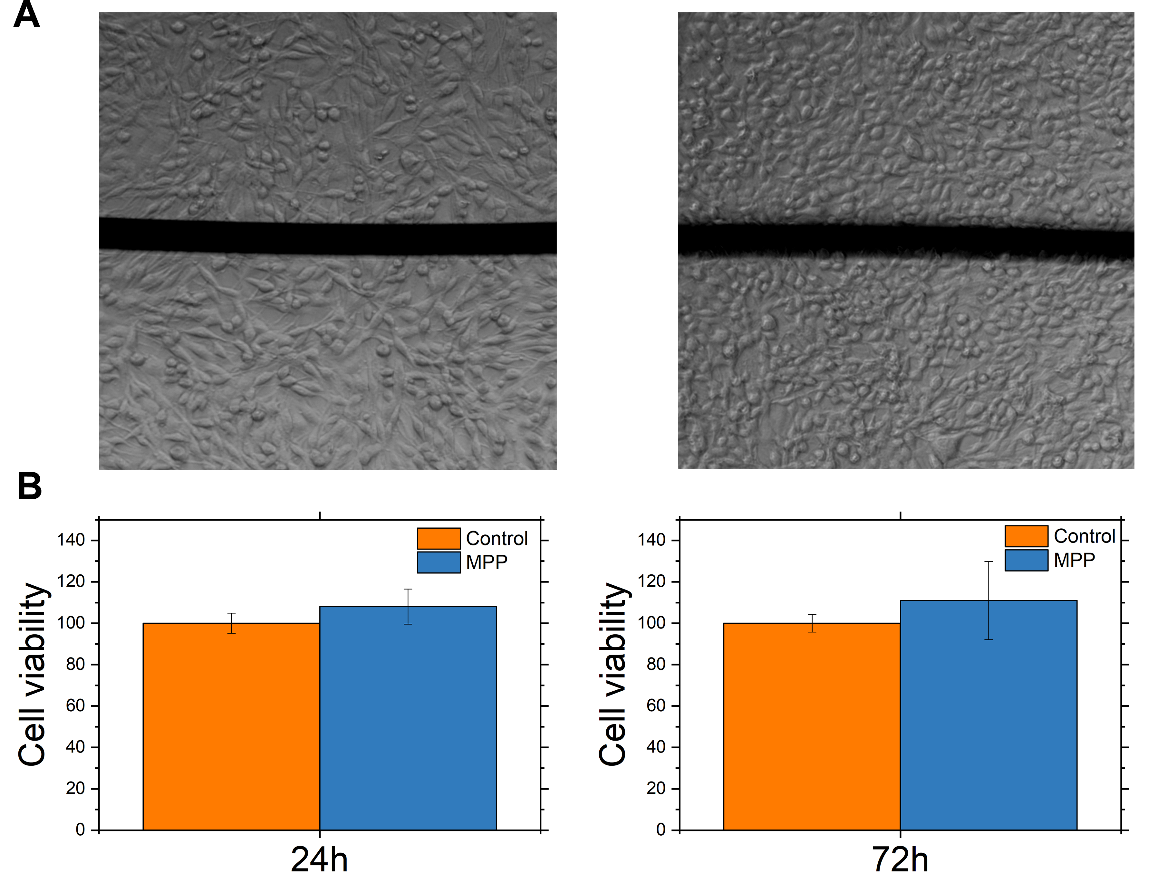

**Fig. S3. PC12 cytotoxicity test.** **(A)** Optical images of cells coexisting with fibers after 24 hours (left) and 72 hours (right). **(B)** Cell viability.

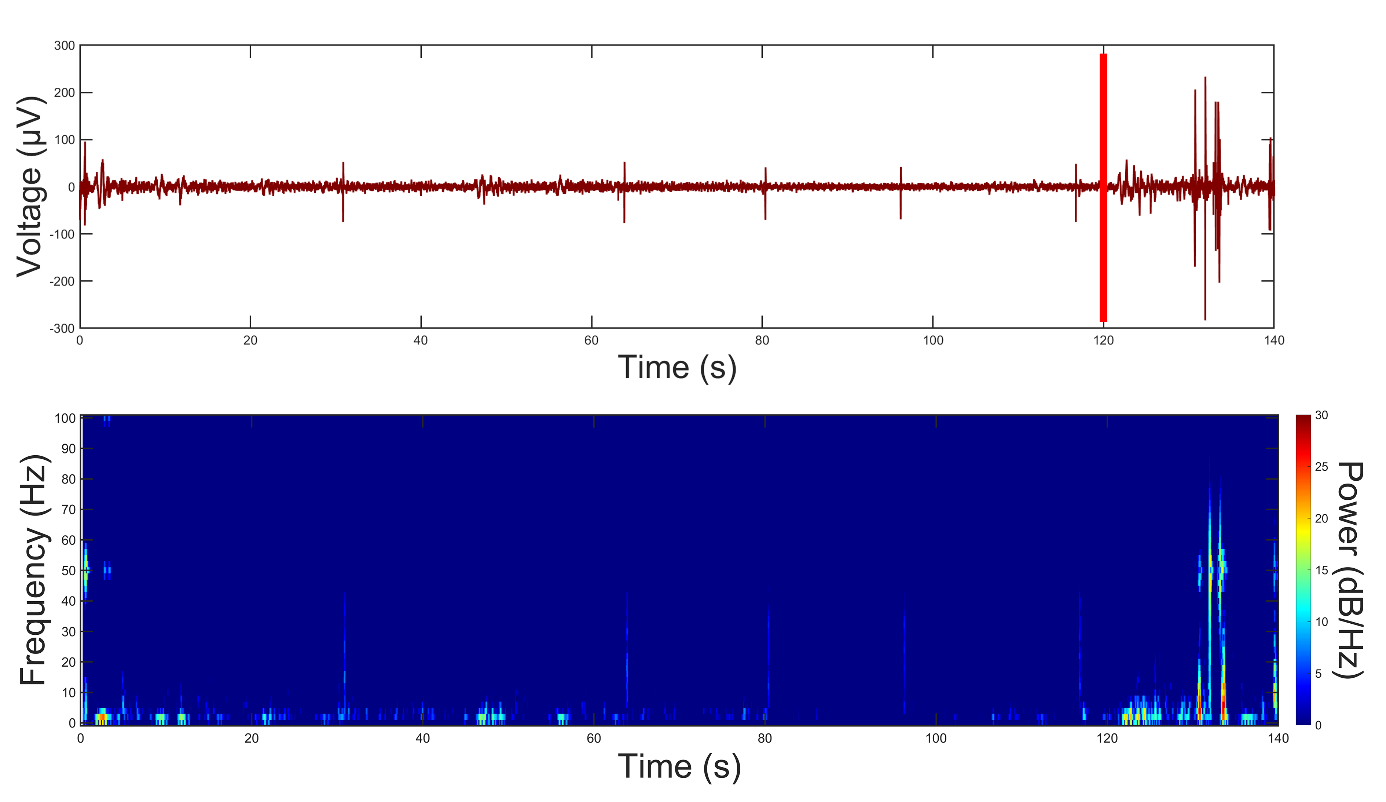

**Fig. S4. Skin electrical signals caused by sympathetic nerve excitation.** The recordings and spectrogram before and after sympathetic nerve excitation. The red line indicates that the alarm goes off.
